# Deep Mediation Analysis for Multimodal Genotype-Imaging Associations with Disease Phenotypes

**DOI:** 10.64898/2025.11.20.689376

**Authors:** Vahid Golderzahi, Guan-Jie Wang, Jacob Hu, Chen-Hsiang Yeang

**Affiliations:** Institute of Statistical Science, Academia Sinica, Taipei, Taiwan

**Keywords:** Multimodal Associations, Genome Wide Association Study, Mediation Analysis, Convolutional Neural Network, Feature Extraction, Alzheimer’s Disease

## Abstract

Both genotype and imaging data carry entangled information about the phenotypes. Associations between genotypes and phenotypes can be manifested or absent on biomedical images. While there are abundant multimodal association studies integrating genotype and imaging data, few of them disentangle their direct and indirect effects in associations. We propose GIF (Genotype-Image-Phenotype), a novel modeling framework to predict phenotypes by genotype and imaging data with a mediation structure of these three sets of variables. GIF constitutes a backbone of mediation path genotype→image→phenotype, and direct genotype→phenotype and image→phenotype paths capturing the additional direct associations not explained by the mediation model. To implement the mediation model with a convolutional neural network (CNN), we constructed a CNN to predict genotypes with images and then employed the joint embedding vectors of the CNN to predict phenotypes. The direct links were sequentially augmented to the mediation model to further improve prediction accuracy. We validated GIF on three datasets: (1) a synthetic polygon dataset where the presence or absence of a polygon specie indicates a genotype and selected combinations of multiple polygon species indicate phenotypes, (2) the PASCAL VOL dataset of object detection and action recognition, where the presence or absence of an object class indicates a genotype and the presence or absence of an action class indicates a phenotype, (3) the ADNI dataset of Alzheimer’s disease diagnosis comprising the genotype and imaging data of subjects with three cognitive states. GIF prediction outcomes on the ADNI data indicate the associations between genotypes and phenotypes are primarily mediated by image features, and images provide additional information about phenotypes which is not attributed to genotype variations.

## 1 Introduction

Emerging AI technologies enable significant advance in discerning phenotypes from large-scale genomic and high-dimensional medical imaging data [1–3]. Associations with the unimodal data of genotypes (Genome-wide association study or GWAS) and images have shown promising results and high impacts on disease diagnosis [4–7]. However, multimodal data integration algorithms yield even greater performances than uni-modal models [8–12]. Multimodal integration methods can be subdivided into three categories in terms of the entry points of integration in the models [10, 13, 14]. Early integration concatenates heterogeneous features before feeding to the model. Some methods extract low-dimensional features from genetic and image data independently and concatenate the multimodal features [15]. Others employ more involved feature merging algorithms such as the high-order graph matching-based feature selection (HGM-FS) to generate the joint features as the model inputs [16]. Intermediate integration learns modality-specific embeddings with deep neural networks (DNNs) separately and concatenates them in a joint DNN. A common approach of fusing modality-specific embeddings is to employ the attention-based transformer models to capture interactions between modalities [8, 17, 18]. Others apply contrastive learning to encode genetic and image data into the joint latent space [12]. Late integration aggregates the model outputs from multiple modes, such as an early-late fusion (ELF) approach [13]. The choice of the fusion strategy (early, intermediate, or late) highly depends on datasets [13, 19, 20].

While some data integration methods are highly effective, they do not explicitly capture the relations of data modes and phenotypes. A simple and intuitive relation between genotypes, images and phenotypes is the existence of three causal/association paths: (1) the effects of genotypes on phenotypes are manifested in images, (2) the effects of genotypes on phenotypes do not appear in images, (3) some determinants of phenotypes are non-genetic and manifested in images. This relation is captured by mediation analysis, as image features often serve as potential mediators between genetic risk and clinical expression [21,22]. While classical mediation analysis handles low-dimensional data with linear models [23], the genotype-image fusion handles high-dimensional data with nonlinear models. Several approaches are proposed to resolve this dilemma. Some methods predict phenotypes by images using autoencoders or image classification models, extract image features from the embedding vectors, and then perform associations between the multi-dimensional genotype features and image features [24–26]. The high-dimensional SNP data are often reduced to low-dimensional features by auto-encoders or principal component analysis. Alternatively, rather than inferring the image features by DNN models, other methods use the annotated image-derived phenotypes (IDP) as low-dimensional image features and perform mediation analysis [27, 28]. In addition, one study first performs genotype-image association by feeding the genotype features to a generative adversarial network (GAN) for images, and then uses the trained genotype features to fit the phenotype [29].

Despite their utility, prior studies on genotype/image mediation analysis have two limitations. They focus primarily on the indirect path of genotype → image → phenotype but do not disentangle its effects from the two direct paths genotype → phenotype and image → phenotype. Most of them perform genotype → image association with surrogate variables of genotypes (e.g., PCA projections of genotype features) and images (e.g., IDPs) rather than directly operating on the original genotype and image data. To address these shortcomings, we propose a mediation analysis based neural network model to integrate genotype and image data for phenotype association, and term this model GIF (Genotype - Image - Phenotype). GIF explicitly constructs the models of the three association paths and iteratively estimates the model parameters of each path conditioned on the models inferred at the previous iterations, with the order of (1) the indirect path, (2) the direct genotype → phenotype path, (3) the direct image → phenotype path. The mediation model is established by first fitting the genotype labels with images using a convolutional neural network (CNN) model, and then employing the embedding vectors of the trained CNN to fit the phenotype labels. We validated GIF on three datasets: (1) a synthetic dataset of polygon images, (2) the PASCAL VOL dataset [30] of object detection and action recognition, (3) the ADNI dataset [31] of Alzheimer’s disease diagnosis. Analysis results on these datasets demonstrate the capacity of GIF to disentangle the effects of direct and indirect association paths and predict phenotypes.

## 2 Methodology

The GIF model of genotype/image associations with phenotypes is illustrated in the schematic diagram in Fig. 1. There are three sets of variables: *G* – genotypes, *I* – images, and *Φ* – phenotypes. The associations of *Φ* with *G* and *I* are realized by three paths. The red links represent the indirect model where the *G* → *Φ* associations are mediated by *I*. The blue and green links represent the direct models where the *G* → *Φ* and *I* → *Φ* associations are not explained by the mediation model. While this architecture resembles the mediation analysis [23], GIF differs from standard mediation analysis in three aspects. First, we leveraged the power of deep neural networks (DNNs) to predict class labels by image data, rather than the regression models commonly used in mediation analysis. Second, images are high dimensional data which are difficult to handle in standard mediation analysis. We developed a novel algorithm to perform mediation by linking the image features with both genotypes and phenotypes. Third, there are two *I* → *Φ* links, one indicates the image-phenotype associations attributed to the genotypes, and the other indicates the image-phenotype associations independent of genotypes.

**Fig 1.**
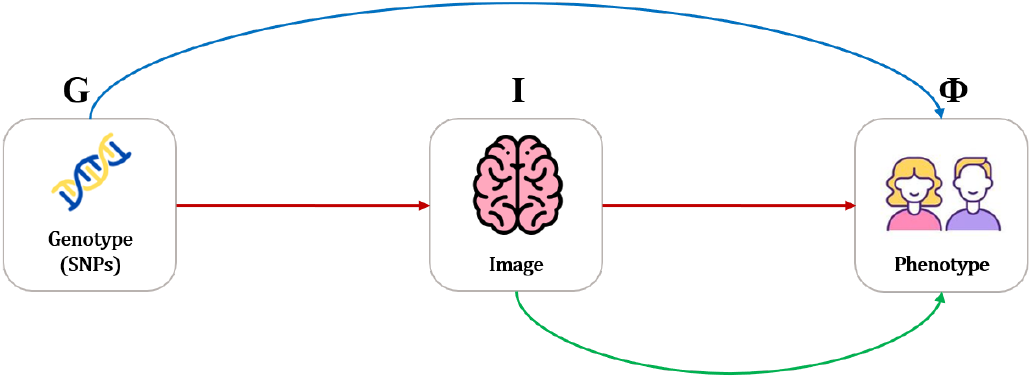
An overview of the proposed GIF model.

The backbone of GIF is a mediation model (*G* → *I* → *Φ*) using DNNs to perform associations. A mediation model denotes existence of two types of models capturing the dependency of genotype and phenotype variables with the same set of variables (features) in images. While variables in *G* and *Φ* are well defined, variables in *I* are elusive. The raw 2D image data can be treated as *d* ≡ *d_x_* × *d_y_* dimensional vectors where *d_x_* and *d_y_* denote the numbers of pixels along *x* and *y* axes, yet this representation often yields poor predictions/associations with the output variables. Instead, we passed an image into a DNN model and treated the neuron outputs of the second last layer as the embedding vector (or features) of the image. In the present study, we chose a pretrained convolutional neural network ResNet-50 [32] as a DNN model, yet other network architectures of image data such as auto encoder/decoders [33] and vision transformers [34] can also be employed in the GIF framework.

The image embedding vectors should simultaneously carry the information about *G* → *I* and *I* → *Φ* associations. All the DNN models predicting the phenotype labels with image data realize the *I* → *Φ* association. In contrast, the *G* → *I* association is difficult to implement because the input variables (genotypes) are discrete and the output variables (image features) are continuous and high-dimensional. Alternatively, we reverted the association direction and perform *I* → *G* association of selected SNPs with CNN, then applied the image features of the model to predict the phenotype (*I* → *Φ* association). Mediation was achieved as the same image features were used to fit both genotypes and phenotypes. On top of the mediation model, we sequentially augmented *G* → *Φ* and *I* → *Φ* links that provided additional power to predict the phenotype labels.

### 2.1 Training the GIF model

Fig. 2 illustrates the procedure of training the GIF model. It consists of the following sequential steps. At step 1, we trained the ResNet-50 model to establish the *I* → *G* links. Rather than including the entire genotype data containing millions of SNPs, we selected a small number of SNPs which have strong or known associations with the phenotype. Each SNP possesses three possible discrete values 0, 1, 2 counting the number of minor alleles. For each SNP, we compressed the allele values to binary labels by collapsing 1 and 2 into class 1 due to the rarity of homozygote minor alleles (2) in the population. The outputs of the second last layer of ResNet-50 are a 2048-dimensional embedding vector. We added a fully connected layer to reduce the dimension of the embedding vector from 2048 to 16. Finally, a fully-connected layer connects the 16-dimensional embedding vector to the joint outputs of the genotype class labels (containing 2 × *K* neurons where *K* is the number of selected SNPs in *G*). A multi-class, multi-label cross entropy loss function was employed to quantify the match of predicted and true class labels over selected SNPs:

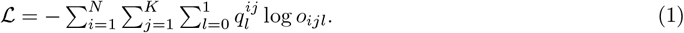

where *i, j, l* denote indices of instances, SNPs, and binary labels, 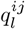 the one-hot probability derived from the class label of instance *i* and SNP *j* (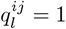 if the reported class label is *l* and 0 otherwise), and *o_ijl_* the normalized model output of instance *i* on the neuron corresponding to SNP *j* and class *l*. Starting with the ResNet-50 default parameter values as the initial weights, we incurred stochastic gradient descent (SGD) to update the weights to minimize the loss.

At step 2, we trained the *I* → *Φ* links and completed the mediation model. Rather than retraining the ResNet-50 model to predict the phenotype labels with images, we used the joint embedding vector inferred from step 1 to fit the phenotypes. The mediation model comprises a ResNet-50 model with the last fully connected layer connecting the 16-dimensional embedding vector to *f* (the number of phenotype labels) output neurons. The weights up to the second last layer were all frozen to the values trained at step 1. At step 2 we only updated the weights of the last layer to minimize the cross entropy loss of phenotype prediction. This model meets the semantic requirement of the *G* → *I* → *Φ* path because the joint embedding vectors in *I* are simultaneously associated with *G* and *Φ*.

At step 3, we trained the direct *G* → *Φ* links conditioned on the mediation model. We selected a logistic regression model to capture *G* → *Φ* dependency since *G* is a low-dimensional and discrete data. Given a *K*-dimensional genotype data **X** and *f* phenotype labels, the genotype feature **Y** is a *f* -dimensional vector linearly transformed from **X**: **Y** = ***β*** · **X. Y** joins the embedding vectors of step 2 to connect to the *f* output neurons of phenotype labels. The weights of all links in the mediation model were fixed, and only the ***β*** coefficients and the weights from **Y** to *Φ* (the dark blue links in Fig. 2) were updated by SGD. This model captures the additional genotype-phenotype association conditioned on the mediation model, hence meets the semantic requirement of the *G* → *Φ* path.

Step 4 is optional as it is incurred only when some selected SNPs do not possess strong *I* → *G* associations. The strength of *I* → *G* associations is defined by the prediction accuracy of the *I* → *G* model trained at step 1. If a SNP label is poorly predicted by images, then it does not appear in the mediation model. We termed these SNPs *G*^′^. At step 4, we trained the direct *G*^′^ → *Φ* links conditioned on the mediation model and the *G* → *Φ* links with the same procedure as step 3.

At step 5, we trained the direct *I* → *Φ* links conditioned on the mediation model, *G* → *Φ* and *G*^′^ → *Φ* links. We constructed another ResNet-50 model and connected its 16-dimensional embedding vector to the phenotype neurons at the final fully connected layer. The weights of the newly augmented model were updated by SGD, while the weights of all other existing models were fixed.

**Fig 2.**
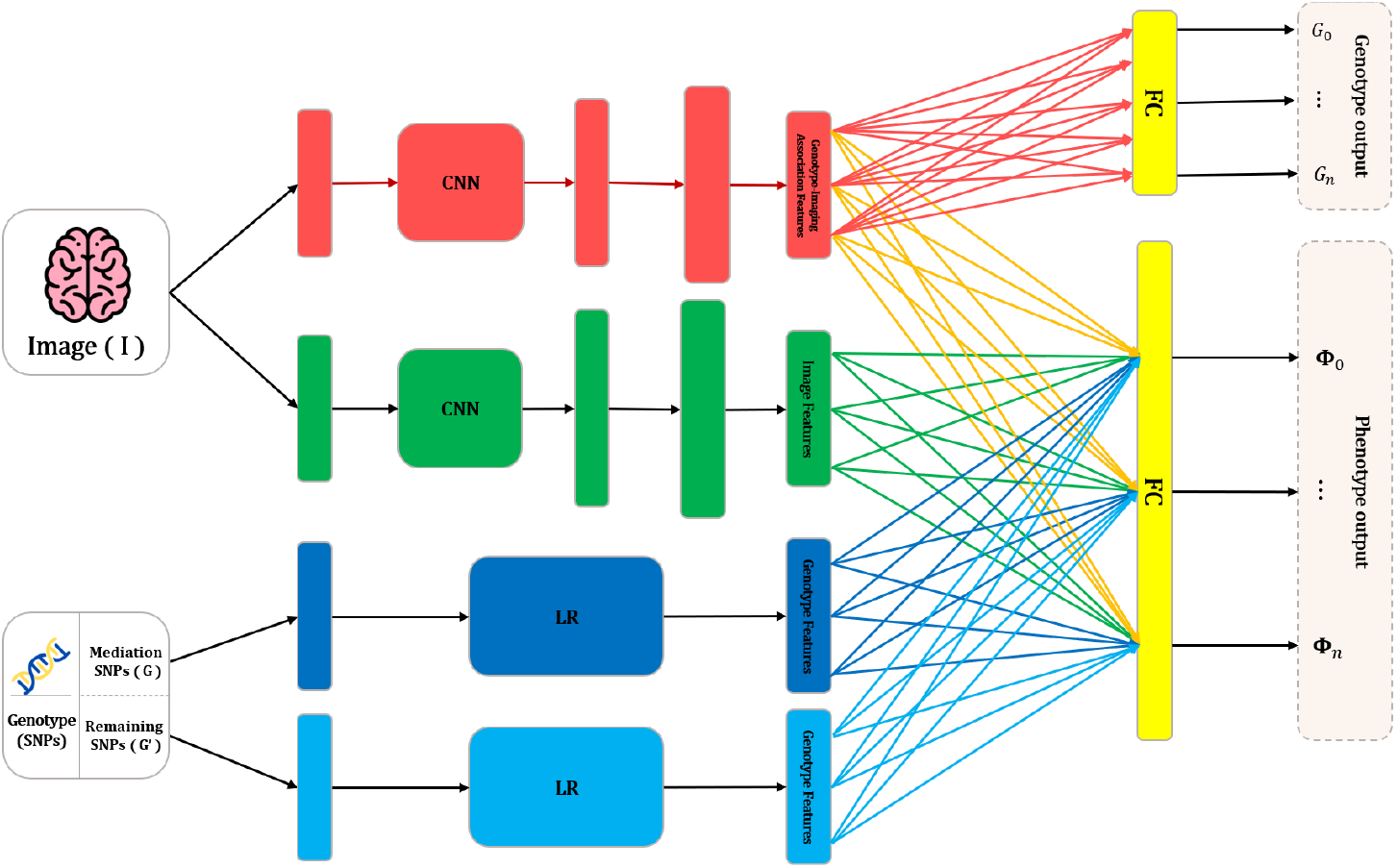
The proposed GIF model architecture. The model weights were sequentially updated with the following order: (1) red, (2) yellow, (3) dark blue, (4) light blue, (5) green.

### 2.2 Data description and processing

We validated the GIF model on the following three datasets.

- We generated a synthetic dataset of images containing multiple species of polygons and their annotations. The dataset consists of 25,000 images, each possesses polygons of four types (“Triangle”, “Square”, “Circle”, “Pentagon”). The *genotype* of an instance is a 1 × 4 binary vector *G* indicating the presence/absence of a polygon type (so there are 4 SNPs in the data). To imitate the direct *G* → *Φ* dependency, we created an auxiliary binary variable *x* independent of image *I* and appended it to the genotype vector *G*. To imitate the direct *I* → *Φ* dependency, we created another polygon type (“Cross”) which may appear in the images but was absent in the genotype data. The phenotype *Φ* is a binary scalar indicating the presence/absence of certain combinations of polygon species and the auxiliary variable. We varied the definition of *Φ* to test the capacity of GIF to capture the three dependency paths, and will discuss these scenarios in the next section.
- The PASCAL VOC 2012 [30] consists of 11,530 annotated images for object detection and action classification. Each image contains objects belonging to 20 classes (e.g., cars, humans, dogs, computers, etc.), and a subset of images are also annotated with 10 (independent) action labels (e.g., horse riding, computer typing, etc). We treated the genotype *G* as the 1 × 20 binary vector indicating presence/absence of each object type, and the phenotype *Φ* as the 1 × 10 binary vector indicating presence/absence of each action type.
- The Alzheimer’s Disease Neuroimaging Initiative (ADNI) [31] is a groundbreaking research project established to study the progression of Alzheimer’s disease through the integration of neuroimaging, clinical assessments, and biomarker data. We solicited the ADNI data of 474 participants including their genotype data of about 3 million SNPs per patient [8], the 2D brain MRI images on the sagittal, coronal, axial planes, and phenotypes of three cognitive categories (Alzheimer’s disease AD, mild cognitive impairment MCI, and control normal CN).

Here we describe the processing of the ADNI dataset. We restricted the datasets to participants with the complete data of genotype, image, and phenotype modalities. There are 474 participants spanning four ADNI studies (ADNI1, ADNI2, ADNI GO, ADNI3), including 45, 158, and 271 patients of AD, MCI and CN categories. If a patient has multiple visits data within a period of disease progression, then we only kept the image and phenotype with the final diagnosis. In addition, we applied filtering methods such as quality control (*QC <* 20), missing rate (*rate >* 0.05), and minor allele frequency (MAF) (*MAF <* 0.01) to capture the most informative and high-quality SNPs. We calculated the odds ratio (OR), Fisher p-value, and FDR based on the Benjamini-Yekutieli (BY) method [35, 36] for each SNP. There were 53 SNPs that passed these filtering criteria, were significantly associated with AD (*p* − *value <* 0.05), and appeared in the GWAS Catalog [37]. Furthermore, we measured the Linkage Diseqilibrium (LD) score for these 53 SNPs and plotted a direct LD graph to illustrate the most correlated SNPs with an *R*^2^ ≥ 0.9 threshold. By clustering SNPs within LD groups, we identified 40 SNP groups and selected one representative SNP for each group.

## 3 Results

We validated the GIF model by applying it to the aforementioned three datasets. On the synthetic polygon data, GIF captures various types of genotype/image dependency with phenotype. On PASCAL VOC 2012, GIF captures the relationship between object classes and action labels of images. On the ADNI dataset, GIF outperforms several other multimodal integration methods in predicting disease phenotypes, and more importantly, demonstrates the associations between genotypes and phenotypes are mediated by image features, and the associations between images and phenotypes are only partially attributed to genotypes.

### 3.1 Association analysis on the synthetic polygon dataset

In this dataset, the genotypes are the presence/absence of each polygon species, and the phenotype is a binary indicator of designed combinations of genotype and image features.

The model architecture for analyzing this dataset slightly differs from that of Fig. 2 in three aspects: (1) A simpler CNN model AlexNet [38] rather than ResNet-50 was employed, (2) The *G*^′^ → *I* association path was absent, (3) In the last layer, we constructed two-dimensional embedding vectors *Z^E^*, *Z^I^* and *Z^G^* for the association models of the three paths *G* → *I* → *Φ, I* → *Φ*, and *G* → *Φ*, and connected the embedding vector neurons of one phenotype value to the corresponding output neuron of the same phenotype value (rather than connecting all embedding vector neurons to all output neurons). We considered six models with varying combinations of the three paths: *M_1_*: *G* → *I* → *Φ, M*_2_: *G* → *Φ, M*_3_: *I* → *Φ, M*_4_: *M*_1_ + *M*_2_, *M*_5_: *M*_1_ + *M*_3_, *M*_6_: *M*_1_ + *M*_2_ + *M*_3_. The polygon dataset was divided into training, validation, and test sets in a 70:10:20 ratio. We trained the model using 20 epochs, a batch size of 256, and a cross-entropy loss function.

To test the capability of GIF to capture various types of genotype/image dependency with phenotype, we constructed four phenotype functions according to genotype and image features. *Φ*_1_, *Φ*_2_, *Φ*_3_, *Φ*_4_ possess the dependency structure of models *M*_1_, *M*_4_, *M*_5_, *M*_6_ respectively.

- *Φ*_1_ = 1 if an image contains triangles and squares, and 0 otherwise.
- *Φ*_2_ = 1 if an image contains triangles, squares and the auxiliary variable *x* = 1, and 0 otherwise.
- *Φ*_3_ = 1 if an image contains triangles, squares, and another specie of polygons (crosses) which does not appear on genotypes, and 0 otherwise.
- *Φ*_4_ = 1 if an image contains triangles, squares, and crosses, and the auxiliary variable *x* = 1, and 0 otherwise.

Table 1 reports the F1-scores of predicting four phenotype functions using six models on the syntehtic polygon data. As expected, each phenotype function is accurately predicted by the corresponding model or more complex models. *Φ*_1_ depends on presence of both triangle and square polygon species that appear in both *G* and *I*, hence is accurately predicted by all models. *Φ*_2_ depends on presence of both triangle and square polygon species and the auxiliary variable *x* which appears in *G* but not in *I*, hence is accurately predicted by *M*_2_, *M*_4_ and *M*_6_. *Φ*_3_ depends on presence of both triangle and square polygon species and the extra polygon type which appears in *I* but not in *G*, hence is accurately predicted by *M*_3_, *M*_5_ and *M*_6_. *Φ*_4_ depends on all the features in *G* and *I*, hence is accurately predicted by only *M*_6_.

**Table 1.**
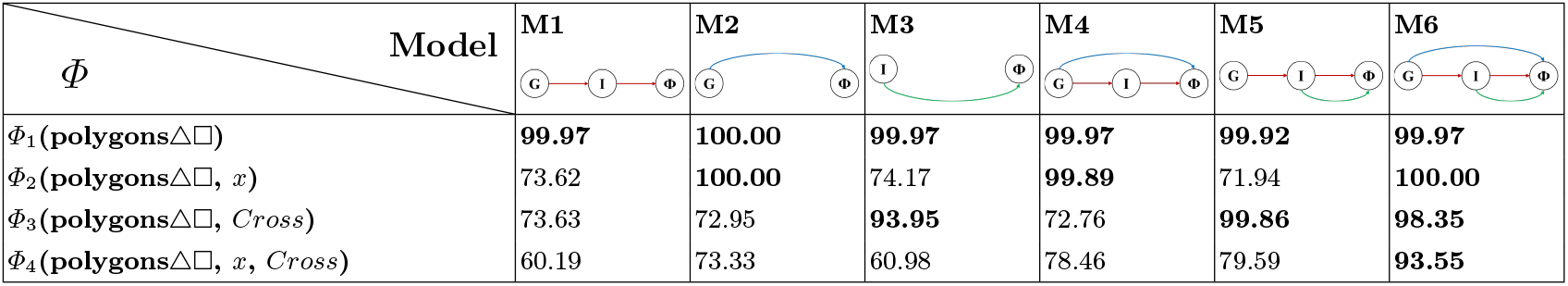
F1-scores (in percentage) of prediction accuracy for four phenotype functions and six models in the polygon dataset.

| $\Phi$ \ Model | M1 | M2 | M3 | M4 | M5 | M6 |
| --- | --- | --- | --- | --- | --- | --- |
| $\Phi_1(\text{polygons}\triangle\square)$ | **99.97** | **100.00** | **99.97** | **99.97** | **99.92** | **99.97** |
| $\Phi_2(\text{polygons}\triangle\square, x)$ | 73.62 | **100.00** | 74.17 | **99.89** | 71.94 | **100.00** |
| $\Phi_3(\text{polygons}\triangle\square, \text{Cross})$ | 73.63 | 72.95 | **93.95** | 72.76 | **99.86** | **98.35** |
| $\Phi_4(\text{polygons}\triangle\square, x, \text{Cross})$ | 60.19 | 73.33 | 60.98 | 78.46 | 79.59 | **93.55** |

### 3.2 Association analysis on the PASCAL VOC dataset

In this dataset, the genotypes are the presence/absence of 20 object classes, and the phenotypes are the presence/absence of 10 action classes. The 20 object classes possess a cluster structure rather than being independent of each other. Hence we adopted two approaches to perform *I* → *G* associations. In the first approach, we treated the 20 object classes as independent genotype variables and predicted 20 genotype labels with images. It is a multi-label classification task utilizing the faster-RCNN architecture with a pretrained ResNet-50 backbone [39]. The training procedure is identical to that of the synthetic polygon data. In the second approach, we classified the 20 objects into several categories sharing similar features (such as animals, furniture, and vehicles). The model has the same ResNet-50 architecture but employs a customized loss function (termed incremental CELoss) to allow incremental clustering of object classes. The dataset was divided into training, validation, and test sets in a 70:10:20 ratio. We trained the model using 15 epochs and a batch size of 32 in all experiments.

In the second approach, we reduced the high-dimensional genotype vectors into the lower dimensional pseudo-class vectors where the genotypes (object classes) were reduced into a small number of categories. We incurred an incremental procedure to learn the categories from the modified CNN architecture and customized loss function. Initially, we replaced the output layer of the CNN model with two neurons *Z*_1_ and *Z*_2_ aiming to collapsing the multi-dimensional object class labels into a one-dimensional binary category labels. The unknown assignment of object classes to each category was implicitly encoded in a customized loss function *CELoss*:

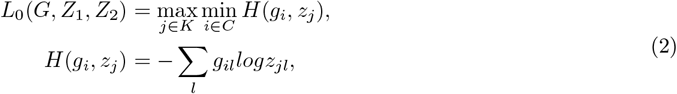

where *K* and *C* are the sets of object classes and hidden binary categories respectively, *g_il_* is the one-hot representation of the object class *i* label in instance *l*, and *z_jl_* is the normalized output value of neuron *Z_j_* for instance *l*. Cross entropy *H*(*g_i_*, *z_j_*) quantifies the match between object class *i* and hidden binary category *j*. The max-min loss function attempts to optimize the match of the worst object class assignment for the best hidden category. By counting the number of instances with each combination of object class labels and hidden category labels, we assigned each object class to either one category (when the counts were concentrated on one category) or no category (otherwise).

In the next steps, we fixed the assignments of object classes to the categories at the previous step (e.g., *Z*_1_ and *Z*_2_ for *L*_0_(*G, Z*_1_, *Z*_2_)), and created a new category neuron to attempt assigning the remaining object classes to it. The incremental *CELoss* at step *K* is:

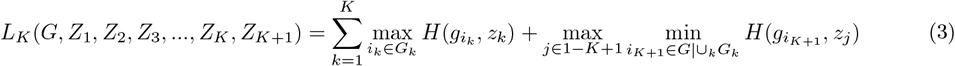

where *G_k_* denotes the object classes assigned to *Z_k_*(*k* ≤ *K*). *L_K_* fixed the object class → hidden category assignments obtained from the previous steps and optimized the assignments of the remaining object classes (*G* |∪ *_k_ G_k_*) to *Z*_1_ − *Z*_*K*+1_. In the present study, we stopped at four hidden categories (*K* = 4) for the PASCAL VOC data. The subsequent steps of the training procedure follow the description in the Methods section. Each of the 10 action classes was treated as a binary phenotype.

Tables 2 and 3 report the prediction results of 10 actions (clinical phenotypes) with 7 models by using the two approaches for *G* → *I* associations, respectively. The two approaches yielded similar prediction outcomes (mean average precision or mAP). *M*_6_ (combination of three association paths) yielded the best or top-ranking prediction outcomes in 8 of 10 action classes, suggesting integration of the three paths gives rise to the most powerful prediction model. In addition, *M*_2_ (marginal association with *G*) and *M*_4_ (direct and indirect associations with *G*) yielded the best or top-ranking performance in the actions involved with the objects which belong to the object classes (Riding-Bike, Riding-Horse) or not (Playing-Instruments, Taking-Photo). In the former case, presence/absence of objects such as bicyles (genotypes) is strongly indicative about the action such as Riding-Bike (phenotype), and images provide redundant information. In the latter case, the objects involved in the action (music instruments and cameras) are not in the 20 object classes, but are probably accompanied with other objects in the 20 object classes (such as co-occurrence of chair and music instruments in instrument playing). Hence genotypes still provide dominant information about phenotypes. For most other actions, both object class labels and images provide essential information. Therefore, the complete GIF model *M*_6_ and the standard integration model *M*_7_ substantially outperform other models.

**Table 2.** Mean avergae precision mAPs (in percentage) of prediction accuracy for ten action classes and seven models in the PASCAL VOC data. The 20 object classes were used as output neurons in the *I* → *G* model of mediation analysis (approach 1).

| Action Classes \ Model | M1 | M2 | M3 | M4 | M5 | M6 | M7 |
| --- | --- | --- | --- | --- | --- | --- | --- |
| Phoning | 81.23 | 81.71 | 81.51 | 81.51 | 81.51 | 81.51 | **82.50** |
| Playing-Instrument | 77.23 | **82.83** | 75.03 | 77.22 | 82.50 | 80.00 | 81.71 |
| Reading | 78.72 | 80.14 | 75.63 | 79.22 | 75.35 | **82.93** | 81.50 |
| Riding-Bike | 84.02 | 94.18 | 75.72 | **95.52** | 76.41 | 93.84 | **95.52** |
| Riding-Horse | 89.81 | **97.85** | 77.34 | **97.85** | 89.53 | 97.00 | 97.00 |
| Running | 82.23 | 84.15 | 76.14 | 83.35 | 80.82 | **85.91** | 84.82 |
| Taking-Photo | 75.43 | **81.72** | 74.63 | 81.33 | 75.00 | 76.15 | 79.62 |
| Using-Computer | 89.11 | 92.18 | 80.63 | 91.82 | 87.61 | **98.12** | 91.82 |
| Walking | 79.35 | 80.53 | 75.25 | 80.52 | 80.11 | **82.73** | 82.01 |
| Jumping | 74.22 | 75.81 | 75.43 | 78.51 | 76.55 | **85.43** | 80.00 |

**Table 3.** Mean avergae precision mAPs (in percentage) of prediction accuracy for ten action classes and seven models in the PASCAL VOC data. The 4 reduced object categories were used as output neurons in the *I* → *G* model of mediation analysis (approach 2).

| Action Classes \ Model | M1 | M2 | M3 | M4 | M5 | M6 | M7 |
| --- | --- | --- | --- | --- | --- | --- | --- |
| Phoning | 81.75 | 81.73 | 81.55 | 81.75 | 81.75 | 81.75 | **82.50** |
| Playing-Instrument | 75.00 | 82.83 | 75.00 | 84.43 | 78.61 | **85.33** | 81.71 |
| Reading | 75.00 | 80.13 | 75.63 | 80.92 | 75.63 | **81.55** | 81.50 |
| Riding-Bike | 75.35 | 94.15 | 75.72 | 94.83 | 81.93 | 94.41 | **95.52** |
| Riding-Horse | 82.02 | **97.81** | 77.35 | **97.81** | 80.15 | **97.81** | 97.00 |
| Running | 74.35 | 84.15 | 76.11 | 83.05 | 79.00 | 83.72 | **84.82** |
| Taking-Photo | 75.41 | **81.73** | 74.60 | 80.62 | 77.85 | 81.33 | 79.62 |
| Using-Computer | 81.22 | 92.15 | 80.63 | **92.94** | 76.25 | **92.94** | 91.82 |
| Walking | 78.60 | 80.53 | 75.25 | 78.61 | 77.80 | **82.35** | 82.01 |
| Jumping | 75.82 | 75.82 | 75.41 | 78.83 | 74.21 | **80.48** | 80.00 |

### 3.3 Association analysis on the ADNI dataset

The ADNI dataset includes the genotype data of about 3 million SNPs per patient, brain MRI images on the sagittal, coronal, axial planes, and the cognitive traits of 474 participants. We incurred the following procedure to select the SNPs for mediation model (*G*) and the direct association model (*G*^′^): (1) Selected 40 candidate SNPs with strong associations between genotypes and phenotypes individually. (2) Among the 40 candidate SNPs selected the top 13 SNPs with strong image → genotype associations, and placed them in the mediation model. (3) Iteratively removed redundant SNPs from the mediation model and stopped when the cross validation prediction accuracy dropped significantly. 11 SNPs remained in the mediation model (*G*), and 29 SNPs had direct connections to the phenotype (*G*^′^). We divided the dataset into training, validation, and test sets in a 70:10:20 ratio, and trained the model using 38 epochs, a batch size of 16, and a class-weight balancing technique [40] (the phenotype classes are imbalanced) throughout the experiments.

Table 4 reports the prediction accuracy (F1-scores) of 12 models with varying combinations of the association paths. Several observations are drawn from the results. First, the mediation model of 11 SNPs (*M* 1) has similar performance to the aggregation of the indirect *G* → *I* → *Φ* path and the direct *G* → *Φ* path (*M* 4.1), indicating that the *G* → *Φ* associations of the 11 SNPs are mediated by the image features. Second, redundance of the direct *G* → *Φ* path of the 11 SNPs is also evident by comparing models *M* 4.2 and *M* 4.3, as well as between models *M* 6.2 and *M* 6.3. Third, the direct *I* → *Φ* path provides considerable extra information by comparing models *M* 4.1 and *M* 6.1 and models *M* 6.2 and *M* 6.3, suggesting the presence of image features which are associated with the Alzheimer’s disease phenotype but not with the 11 selected SNPs. Fourth, the best GIF models (*M* 6.3 and *M* 6.2) are considerably superior to the standard integration model (*M* 7), suggesting that incorporating a possible causal structure can also improve prediction.

**Table 4.** F1-scores (in percentage) of prediction accuracy for the cognitive phenotypes (CN, MCI, AD) and twelve models in the ADNI dataset. Note: *M* 2.1: the direct genotypes-penotypes association with 11 SNPs. *M* 2.2: the direct genotypes-phenotypes association with 40 SNPs.

| Phenotypes \ Model | M1 | M2.1 | M2.2 | M3 | M4.1 | M4.2 |
| --- | --- | --- | --- | --- | --- | --- |
| Prediction (CN, MCI, AD) | 61.35 | 38.07 | 61.29 | 71.19 | 62.14 | 68.46 |
| Phenotypes \ Model | M4.3 | M5 | M6.1 | M6.2 | M6.3 | M7 |
| Prediction (CN, MCI, AD) | 66.98 | 73.91 | 78.22 | 79.06 | **79.73** | 72.98 |

Table 5 compares the prediction performance of the best GIF model *M* 6.3 with three existing methods of genotype-image integration. GIF outperforms Random Forest and MADDi by a large margin in all prediction metrics and outperforms AlzCLIP with a small margin in accuracy.

**Table 5.**
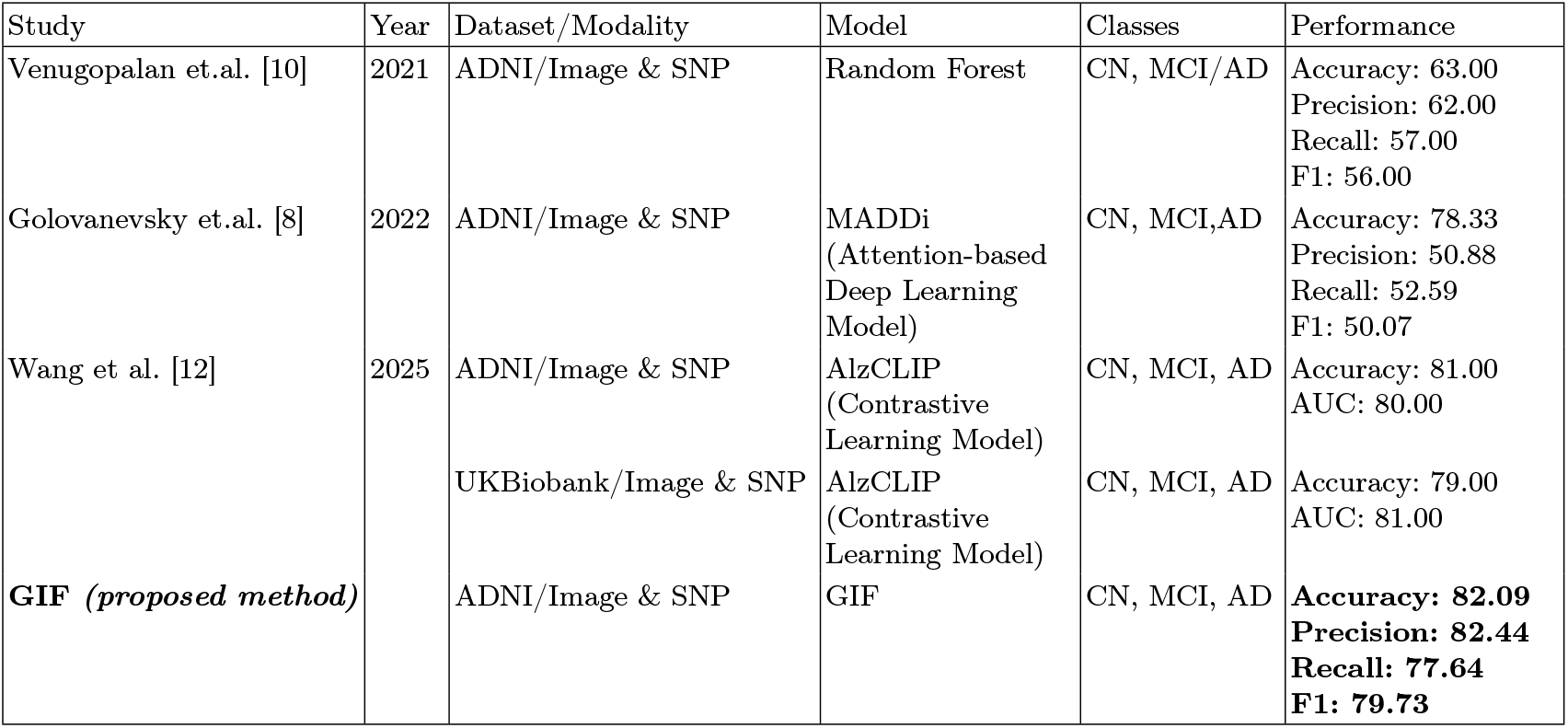
Comparison of the prediction metrics between GIF and three prior studies on the ADNI dataset.

## 4 Conclusion and future work

In this study, we proposed a modeling framework GIF to integrate genotypes and images to predict phenotypes. GIF undertakes mediation analysis by combining the three association paths and implements the association models with deep neural networks. It possesses several unique features absent in the standard multimodal data integration algorithms and mediation analysis as it (1) disentangles the effects of direct and indirect association paths and thus reveals the causal structure of the observables, (2) handles high-dimensional variables and implements nonlinear models in the mediation analysis, (3) allows different association models or DNN architectures to plug into the framework. Analysis on annotated image datasets demonstrates the capacity of GIF to capture the genotype/image/phenotype dependency which is either explicitly designed (the synthetic polygon dataset) or relatively transparent to humans (the PASCAL VOC dataset). Analysis on the ADNI dataset indicates that the genotype → phenotype association of several informative SNPs is mediated by image features, and images provide additional information about phenotypes which is not attributed to genotype variations. Furthermore, GIF yields superior phenotype prediction accuracy rates than several prior data integration methods, suggesting the possible benefits of a causal model over naive integration algorithms in phenotype prediction.

We plan to extend the present work in several directions. First, the prediction outcomes allude the importance of the three association paths, but GIF does not give concrete and interpretable information such as the image features that mediate the phenotype association with specific SNPs and the image features that appear in the direct *I* → *Φ* path. We will enhance GIF to make it capable of retrieving the interpretable features on the association paths. Second, diverse modes of data beyond genotypes and images are available (including electronic health records, biophysical measurements such as EEG and ECG data, and biochemical measurements such as transcriptomic and metabolomic data). We need to extend the simple model (Fig. 1) to handle more complex and diverse causal structures of more diverse modes. Third, the present study serves as a proof-of-concept demonstration of GIF. We plan to further investigate the biomedical and clinical implications of the joint association model in Alzheimer’s disease patients. Fourth, we will also extend the application domains by employing GIF to the multimodal data pertaining to other major diseases such as cancer [41], diabetes [42], and immune diseases [43].

## Source codes and example data

The source codes and example data of running the programs are deposited under github.com/chyeang/GIF.

## Acknowledgments

The authors are supported by Academia Sinica Seed Grant (AS-GCS-113-M05).

